# HDAC6 is a novel regulator of endothelial-to-mesenchymal transition in venous thrombosis

**DOI:** 10.64898/2026.08.27.747544

**Authors:** Marion Pilard, Virginie Gourdou-Latyszenok, Lénaïck Gourhant, Estelle L. Ollivier, Thibaut Riou, Jamal Elhasnaoui, Lanig Civi, Léna Bicrel, Rafael Ricci de Azevedo, Sara Robin, Gilles Pernod, Cécile Tromeur, Francis Couturaud, Catherine A. Lemarié

## Abstract

**Background:** Venous thromboembolism (VTE), which encompasses deep vein thrombosis (DVT) and pulmonary embolism (PE), is a frequent disease associated with thrombus formation and vein wall remodeling. Hence, fibrosis might result from endothelial-to-mesenchymal transition (EndMT), characterized by the loss of endothelial markers and the acquisition of mesenchymal markers. In chronic thromboembolic pulmonary hypertension, transforming growth factor (TGFβ), the most potent inducer of EndMT, impairs thrombus resolution. However, the molecular mechanisms implicated in TGFβ signaling in the context of VTE are unknown. We hypothesized that epigenetic processes regulate the TGFβ signaling pathway in endothelial cells promoting EndMT and vascular fibrosis.

**Aims:** To determine if the histone deacetylase 6 (HDAC6) regulates the TGFβ signaling pathway in endothelial cells promoting EndMT and delays venous thrombosis.

**Methods:** To study the role of HDAC6 in EndMT, endothelial cells were treated with a pharmacological inhibitor (TCS20b) and incubated with TGFβ and thrombin for 2, 3, and 5 days. Real time PCR and Western blot were performed to analyze endothelial and mesenchymal marker expression and TGFβ signaling. An experimental model of VTE was used to study the role of HDAC6 on thrombus size overtime. Animals were treated or not with a specific HDAC6 inhibitor (tubastatin A) for 7 to 21 days. Analysis of RNAseq data sets publicly available were used to confirm our main results. Within group and treatment differences were analyzed using two-way ANOVA and Tukey’s multiple comparisons.

**Results:** Expression of the mesenchymal markers, calponin and transgelin, was increased by TGFβ and thrombin. Interestingly these changes were inhibited in presence of TCS20b. TGFβ mediated these effects through ERK1/2 and HDAC6 activation. Inhibition of HDAC6 *in vivo* reduced thrombus size 7 days after surgery compared to controls. This was associated with reduced expression of the EndMT marker transgelin in endothelial cells compared to the control animals. We found that FN1-EDA expression was associated with EndMT and regulated by HDAC6 *in vitro*. This marker was also associated with thrombosis in the RNAseq data set that we analyzed and potentially in patients with recurrent DVT.

**Conclusion:** We found that HDAC6 regulates EndMT in venous thrombosis and impairs thrombus resolution. HDAC6 also regulates expression FN1-EDA that appears to be a strong marker associated with DVT and DVT recurrence. Thus, HDAC6 might represent an attractive therapeutic target for patients with a high risk of recurrent VTE.

## Introduction

Venous thromboembolism (VTE), which encompasses deep vein thrombosis (DVT) and pulmonary embolism (PE), is a frequent and life-threatening disease. VTE is the third leading cause of cardiovascular death after myocardial infarction and stroke with a mortality rate of 10% at 3 months after PE. VTE affects 1 to 2 per 1000 persons per year and is associated with long-term complications affecting the life expectancy and the quality of life of patients. Main complications of VTE include recurrent (non-fatal and fatal) VTE and long-term sequelae due to incomplete clot resolution in the leg veins (the post-thrombotic syndrome [PTS]) and/or in pulmonary arteries (chronic thromboembolic disease [CTED] and chronic thromboembolic pulmonary hypertension [CTEPH]). Hence, VTE is a major health issue representing 240 million euros per year in France^1^.

One of the major issues is to prevent the recurrence risk, which ranges from 10 to 30% at 5 years, with a case-fatality rate of 10%, depending on whether VTE was provoked by major clinical risk factors (surgery, hormone exposure in women, immobilization, cancer) or not. In more than 50% of cases, VTE occurs in the absence of clinical risk factors (termed “unprovoked” VTE). For these patients, international guidelines recommend treating them with anticoagulant therapy indefinitely to prevent the risk of recurrence^2,3^. However, these treatments are still associated with an important risk of hemorrhage, despite their efficacy at preventing recurrence. Importantly, anticoagulation has little impact on the prevention of chronic sequelae, such as PTS, CTED and CTEPH, which overall affect about one-third of patients^4^. After stopping anticoagulation, recurrence affects about 30% of the patients with unprovoked VTE. Thus, extending indefinitely anticoagulant therapy for all patients with unprovoked VTE is exposing the other 70% patients to an unjustified risk of hemorrhage. Better discriminating which patients with unprovoked VTE have a high risk of recurrent events, is of utmost importance to improve patient care. Currently, no known scores or biomarkers are useful to discriminate patients with high or low risk of recurrence. More importantly, research is needed to decipher pathophysiological mechanisms involved in VTE long-term complications to better prevent evolution of VTE toward a chronic disease.

Endothelial cells play a crucial role in supporting thrombus formation and resolution in collaboration with platelets and leukocytes. It has been long known that thrombus resolution is associated with angiogenesis promoting recanalization of the vessel in association with remodeling mechanisms including fibrosis. However, uncontrolled or extensive fibrosis is an important contributor to the development of pulmonary sequelae such as CTEPH^5^. The underlying mechanisms of fibrosis formation and its contribution to VTE recurrence are still poorly understood. Transforming growth factor-β (TGFβ) is a potent inducer of fibrosis and a cytokine affecting many cellular functions including tissue remodeling and angiogenesis. Importantly, TGFβ is also a master regulator of endothelial-to-mesenchymal transition (EndMT). EndMT is an important pathological process affecting the endothelial phenotype and increasingly involved in cardiovascular diseases. EndMT is characterized by the loss of endothelial markers and the acquisition of several mesenchymal cell markers and properties. TGFβ1 overexpression and enhanced TGFβ1 signaling in endothelial cells has been shown to promote EndMT and contribute to impairment of thrombus resolution^6^. Interestingly, in a mouse model of vein compression, EndMT was found to be both a contributing factor and a consequence of thrombosis, which was associated with loss of anticoagulation and thrombolytic function of endothelial cells^7^.

Epigenetic mechanisms and regulators of post-transcriptional modifications, such as acetylation and methylation, are contributing to epithelial-to-mesenchymal transition (EMT). Because histone deacetylase (HDAC) inhibitors prevent EMT, these proteins have been studied in the context of EndMT. Acetylation is a tightly regulated mechanism by which gene expression and/or protein activity is modulated. Furthermore, HDAC inhibitors have antifibrotic effects in cardiac remodeling by inducing genes repressing cardiac fibroblast activation and extracellular matrix production^8^. HDACs are a family of enzymes divided into 5 categories, class I (HDAC1, −2, −3, −8), class IIa (HDAC4, −5, −7, −9), class IIb (HDAC6, −10), class III (Sirt1–7), and class IV (HDAC11). Among this family, HDAC6 is a structurally and functionally unique member that mainly localized in the cytoplasm. HDAC6 plays a crucial role in the acetylation of non-histone proteins. It can deacetylate substrates, including tubulin, β-catenin, Hsp90 and cortactin. Tubulin acetylation/deacetylation contributes to microtubule-mediated processes such as migration and spreading which might contribute to mesenchymal-like properties acquired by endothelial cells. In addition, HDAC6 expression has been shown to be upregulated by hypoxia^9^. In this study, we investigated the contribution of HDAC6 in TGFβ-induced EndMT in VTE. We sought to characterize EndMT following VTE and how it is regulated downstream of TGFβ.

## Materials and methods

### Animals

Male C57Bl/6J mice from Charles River aged between 8-12 weeks were used for the study. All procedures described were approved by the local animal welfare committee and the French Ministry for Superior Education and Research according to the European Community guidelines.

### Treatment and venous thrombosis model

Venous thrombosis was induced using the electrolytic inferior vena cava (IVC) model (EIM) according to the literature^10^. Briefly, mice were anesthetized by inhalational isoflurane and placed in the supine position on a heating pad set to 37°C. Following a midline laparotomy, a section of the IVC between the renal and left common iliac veins was cleared and separated from the aorta by dissection. All side branches were ligated with 7-0 Prolene suture (Ethicon, F2854H). An anode of 27G stainless-steel needle electrode (NE-115B, Nihon Koden) was inserted into the caudal IVC and attached to the anterior wall. The cathode was introduced subcutaneously. A constant current of 250 μA was applied to the IVC for 15 minutes. The needle was removed, and hemostasis was achieved. The abdominal wall and the skin were closed separately using a 7-0 prolene suture for the muscles and a 6-0 prolene suture for the skin (Ethicon, 6-0). The timeline and experimental groups are illustrated in Figure 6A. Animals were separated in two treatment groups. The control groups received a daily intraperitoneal (IP) injection of vehicle (PEG300- Tween80- DMSO- H_2_O) whereas the experimental groups received 30 mg/kg of tubastatin A (Selleckchem, S8049). Animals were then sacrificed at days 7, 14, 17 and 21 following the EIM model.

### Histology

Thrombi were embedded in OCT and serial 7-μm frozen sections were cut using cryostat and transferred onto gelatin-coated slides. For histology, slides were stained using the Carstairs method to distinguish fibrin, platelet, collagen and red blood cells. Thrombus surface area was quantified by measuring areas on 3 slides per animals and averaged.

### Immunofluorescence

For immunofluorescence staining, samples were incubated with 0.5% Triton X-100 for 10 min followed by 20 % goat or rabbit serum and 5 % BSA for 30 min at room temperature. The sections were incubated for 1h at room temperature in 0.3% BSA in PBS with HDAC6 (1/50; Sigma-Aldrich). After washing, sections were incubated with the 555-nm Alexa Fluor-conjugated secondary antibody in 0.3% BSA in PBS for 1 h at room temperature. Nuclei were labeled with DAPI.

### In situ hybridization RNAscope

Samples were processed according to the manufacturer’s instructions (BioTechne). Slides were prepared for RNAscope through a series of incubation steps including dehydration, peroxide blocking, target retrieval, protease treatment and hybridization with specific Z probes. Briefly, mRNA in the tissue were detected by incubating with RNAscope Z probes for either transgelin (BioTechne catalogue number 480331-C2) or TGFβ (BioTechne catalogue number 407751-C2) for 2h at 40°C. The RNAscope Multiplex Fluorescent Detection Kit v2 (catalogue number 323110) (BioTechne) was used according to the manufacturer’s instructions. Briefly, slides were successively hybridized with AMP1 for 30 min at 40°C, AMP2 for 30 min at 40°C and AMP3 for 15 min at 40°C. Slides were then incubated with TSA vivid 650.

Immediately, after RNAscope, samples were incubated overnight at 4°C with the primary antibody anti-CD31 (1/100; BD Biosciences) diluted in TBS-1% BSA. After washing steps, sections were incubated with the secondary antibody for 1h. The secondary antibody used was: donkey anti-rat AF555 (1/200, Invitrogen). Finally, slides were mounted with Prolong Gold anti-fade reagent with DAPI mounting medium (Invitrogen). A probe specific to the housekeeping genes Polr2a, PPIB, and UBC was used as a positive control and a probe specific to the bacterial dapB gene as a negative control probe (BioTechne catalogue numbers 320881-C1-C2-C3 and 320871-C1-C2-C3, respectively). Images were taken with an axio observer microscope (Zeiss). Images were analyzed with Fiji.

### Patients with venous thromboembolism

Patient samples were selected from two similarly designed double-blind randomized trials where patients with a first unprovoked deep vein thrombosis (DVT) (PADIS-DVT study^11^) and those with a first unprovoked pulmonary embolism (PE) (PADIS-PE study^2^) were included after an initial 6-month anticoagulation and randomized to an additional 18-month warfarin versus placebo. All patients were followed during 24 months study treatment end. In these studies, plasma was collected at acute phase of VTE and at 6 months (while on anticoagulant treatment), at 7 and 24 months (without anticoagulation in the placebo group, while on anticoagulant treatment in the warfarin group), and at 25 and 48 months without anticoagulation in the two groups. Plasma were used to quantify fibronectin-EDA. We selected plasma from the placebo group at 7, 24, 25 and 48 months for the recurrent and non-recurrent groups. Patients’ characteristics are summarized in Table I (Supplementary information).

### RNAseq data processing and analysis

To characterize fibronectin (FN1)-EDA regulation under thrombotic conditions, alternative splicing of the FN1 gene was analyzed using rMATS^12^ and IsoformSwitchAnalyzeR^13^ in two independent publicly available bulk RNAseq datasets generated from murine and porcine models of DVT and PE^14–16^. Particular attention was given to the regulation of the EDA-encoding exon 33 (E33). To enable cross-species comparison, murine and porcine genomic coordinates were converted to the human hg38 reference genome using liftOver^17^. Differences in E33 expression and splicing were assessed by quantifying exon expression levels together with RNA-seq reads supporting exon inclusion and exclusion. Splicing patterns across conditions and species were visualized using Sashimi plots^18^.

### Endothelial cell culture and treatment

HUVECs (Promocell) were grown in 50/50 of endothelial cell basal medium (EBM-2) supplemented with and endothelial cell bullet kit (EGM-2) (Lonza) and of Dulbecco’s Modified Eagle’s Medium F-12 (DMEM/F-12) supplemented with 10% FBS and 1% of penicillin-streptomycin. Cells were used between passage 3 and 5 in tissue culture dishes coated with 0.1% gelatin and maintained at 37°C in a humidified incubator at 5% CO2. For endothelial-to-mesenchymal assay, 4 x 10^5^ HUVECs were plated in 60 mm culture dishes in EBM-2 supplemented with hydrocortisone, ascorbic acid, GA-1000, 10% FBS and TGFβ2 (10 ng/mL) or thrombin (1 U/mL). For the control condition, cells were cultured in the same medium but without TGFβ2 or thrombin. When indicated HDAC6 was inhibited with vorinostat (5 μM, Cayman Chemical Compagny), TCS 20b (5 µM, Tocris) or tubacin A (20 µM, Tocris). HUVECs were treated for 30 minutes, or several days (2, 3 or 5 days) and used for western blot, real-time PCR or immunofluorescence analyses.

### RNA isolation, Reverse-Transcription and quantitative real-time PCR

Gene expression was evaluated in HUVECs by quantitative real-time PCR (qRT-PCR). Total RNA was extracted from cultured cells using a commercial kit following the manufacturer’s instructions (Nagel-Macherey). Four hundred ng to one μg of total RNA were reverse-transcribed as per the manufacturer’s instructions (Ozyme). The SYBRgreen intercalant was used for amplification detection with the Fast SYBRgreen master mix (Ozyme). Primers were designed using the Primer Express Software (Applied Biosystems). The TATA box binding protein (TBP) gene for human cells were used for normalization. Fold changes were calculated using the ΔCt method and results were expressed as fold change ± SEM of five to seven independent experiments. Human primer sequences are listed in Table I.

### Western Blot

HUVECs were lysed with commercially available mammalian protein extraction reagent buffer (Fisher Scientific). Twenty-five µg of protein were separated by 10% sodium dodecyl sulfate polyacrylamide gel electrophoresis (SDS-PAGE) and transferred to nitrocellulose membranes (Fisher Scientific). Western blot analysis was performed with the following antibodies: mouse anti-HDAC6 (1:1000, #sc-28386), mouse anti-GAPDH (1:1000, #sc-32233, Santa Cruz Technology), rabbit anti-alpha-tubulin (1:1000, #2125), rabbit anti-acetyl-alpha-tubulin (1:1000, #5335S), rabbit anti-phospho-ERK1/2 (1:1000, #9101S) and rabbit anti-ERK1/2 (1:1000, #9102S, Cell Signaling). Anti-rabbit or anti-mouse coupled to HRP were used as secondary antibodies (1:2000, Dako). Signals were revealed by chemiluminescence (Fisher Scientific) with the GeneGnome molecular chemiluminescence imager system (Syngene) and quantified by densitometry with Quantity one software (Bio-Rad).

### Statistical analysis

Data are presented as means ± SEM from multiple independent experiments. Normality was assessed using the d’Agostino-Pearson test. When comparing two experimental groups (WT mice with or without tubastatin A treatment), Student’s t test was used. When comparing more than 2 groups, with different treatments (vehicle, TGFβ2, thrombin, TCS 20b or tubacin), two-way ANOVA followed by Tukey’s multiple comparisons test were done. When normality was not present, Kruskal-Wallis test was performed. GraphPad Prism software was used to analyze data and to generate graphs (version 9.0.1). A value of P < 0.05 was considered statistically significant.

## Results

### TGFβ2 and thrombin induce EndMT through HDACs

First, to determine if TGFβ2 and thrombin induce EndMT, we evaluated mesenchymal (transgelin (TGLN) and calponin (CNN1)) and endothelial (VE-cadherin (CDH5)) markers in HUVECs. After 3 days, only TGLN was induced by TGFβ2 and TGFβ2 with thrombin treatment. Next, HUVECs were treated with vorinostat, a HDAC inhibitor. Interestingly, vorinostat blocked TGLN induction following treatment with TGFβ2, thrombin or both (Figure 1A). TGFβ2 and TGFβ2 with thrombin treatment had no effect on CNN1 mRNA expression (Figure 1B). We also evaluated the expression of the endothelial gene CDH5 in the same experimental conditions. TGFβ2, thrombin or vorinostat had no effect on CDH5 mRNA expression (Figure 1C).

**Figure 1.**
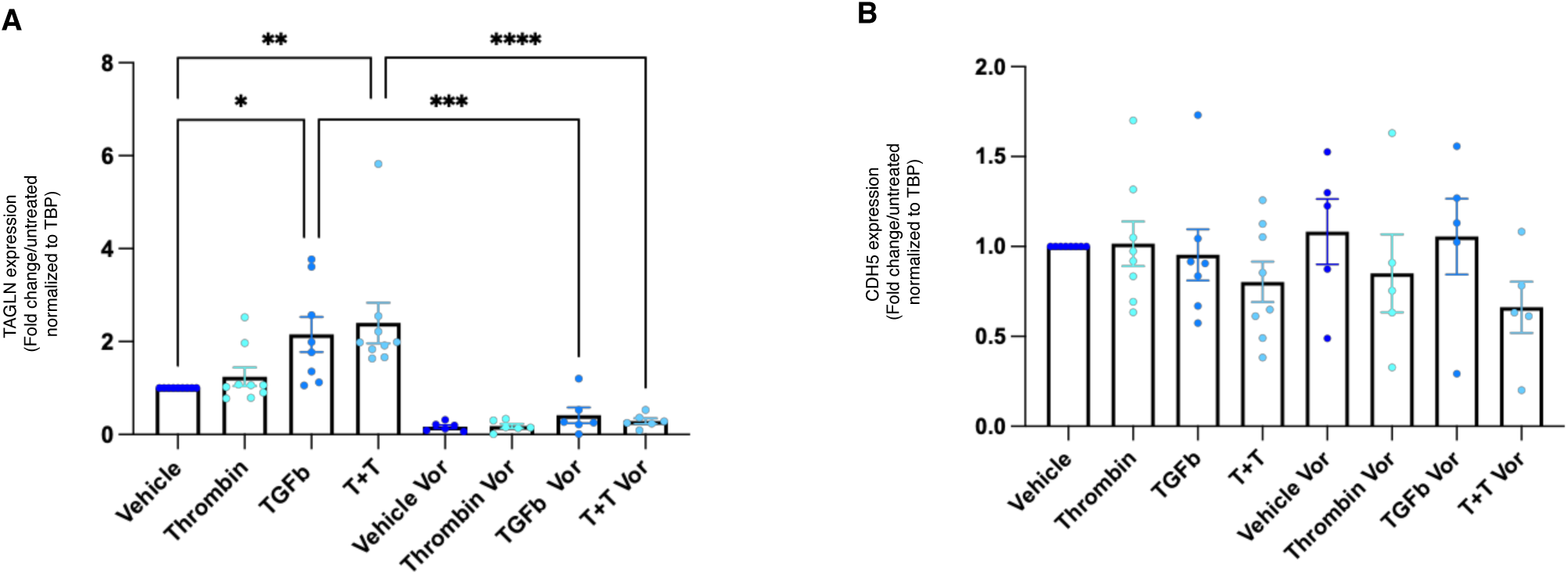
HDACs promote EndMT in response to TGFβ and thrombin. (A) Transgelin (TAGLN) mRNA expression is increased by thrombin or TGFβ2 and further enhanced when cells are stimulated by thrombin and TGFβ2 in combination for 3 days. Interestingly, inhibition of HDACs with vorinostat blocked thrombin and TGFβ2-induced TAGLN mRNA expression. (B) However, neither calponin (CNN1) nor (C) VE-cadherin (CDH5) are induced or regulated by thrombin, TGFβ2 or vorinostat. (N=6; *P<0.05; **P<0.01; ***P<0.001; ****P<0.0001).

### HDAC6 inhibition enhanced TGFβ2-induced alpha-tubulin acetylation

We investigated the impact of TGFβ2 and thrombin on the acetylation status of α-tubulin with or without a selective HDAC6 inhibitor, TSC 20b (Figure 2A). In basal condition, α-tubulin was deacetylated. TGFβ2, alone or in association with thrombin, did not modify the level of acetylation of α-tubulin compared to the vehicle condition. When HDAC6 was inhibited by TSC 20b, the level of acetylated α-tubulin was increased in control and further increased with TGFβ2 (Figure 2Bi). These data confirm that TSC 20b treatment inhibited HDAC6 activity. We tested if TGFβ2 and/or thrombin modified HDAC6 expression. After 5 days of treatment, HDAC6 protein expression was not modified by TGFβ2, thrombin or TCS 20b (Figure 2Bii). Accordingly, HDAC6 mRNA expression was not modified by TGFβ2, thrombin or TSC 20b treatment (Figure 2C).

**Figure 2.**
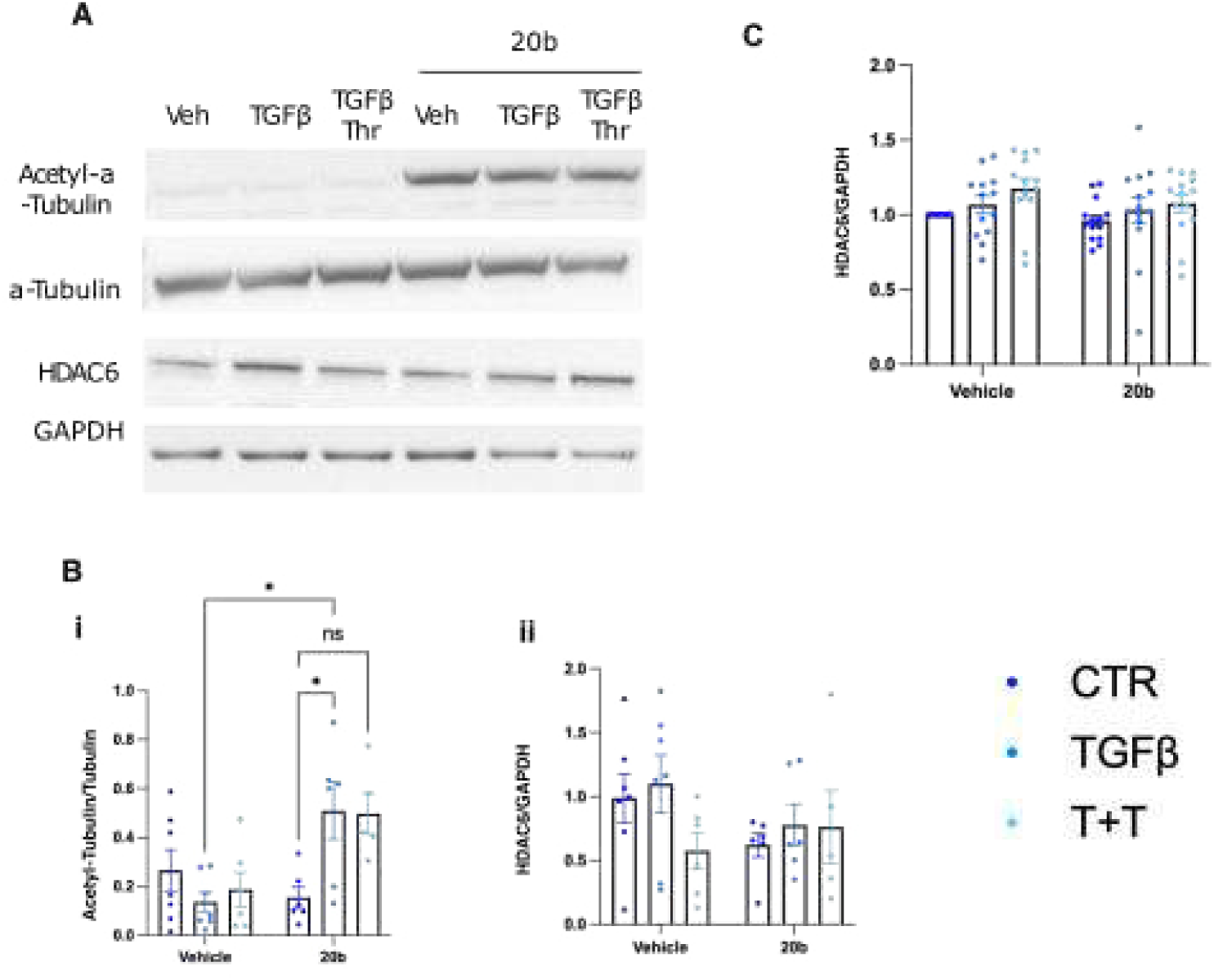
HDAC6 activity induced αtubulin deacetylation. (A) Representative western blot for HDAC6, acetyl-α-tubulin, α-tubulin, HDAC6 and GAPDH in HUVECs treated with TGFβ2 or TGFβ2 and thrombin in presence or not of TCS20b. (B) (i) When HDAC6 is active, α-tubulin is deacetylated. However, when HDAC6 activity is inhibited by TCS20b, α-tubulin is acetylated. ii) Quantifications of HDAC6 protein expression showed that treatment with TGFβ2 or TGFβ2 and thrombin do not modify HDAC6 expression. (C) HDAC6 mRNA expression is not modified by treatment with TGFβ2 alone or TGFβ2 and thrombin. Similarly, treatment with a pharmacological inhibitor of HDAC6 does not affect its expression. (N=6-7; *P<0.05).

### HDAC6 inhibition reduces EndMT-associated gene expression

We evaluated if HDAC6 inhibition regulates expression of genes associated with EndMT after incubation with TGFβ2 with or without thrombin. After 5 days, endothelial cells incubated with TGFβ2 or TGFβ2 and thrombin expressed several mesenchymal markers including TGLN, CNN1 and fibronectin-EDA (FN1-EDA) (Figure 3A-C). In addition, the expression of TGFβ signaling pathway components activin receptor-like kinase 5 (ALK5; TGFβ type I receptor (TGFBR1)) and snail family transcriptional repressor factor 2 (SNAI2) were increased by TGFβ2 or TGFβ2 and thrombin (Figure 3D-E). Interestingly, when HDAC6 is inhibited by TCS 20b, expression of mesenchymal markers and TGFβ signaling pathway components listed above were no longer modified by TGFβ2 or TGFβ2 and thrombin treatment. TGFβ expression was not modified by any of the treatments (Figure 3F). These data suggest that HDAC6 regulates the expression of some mesenchymal markers in response to TGFβ2 or TGFβ2 and thrombin.

**Figure 3.**
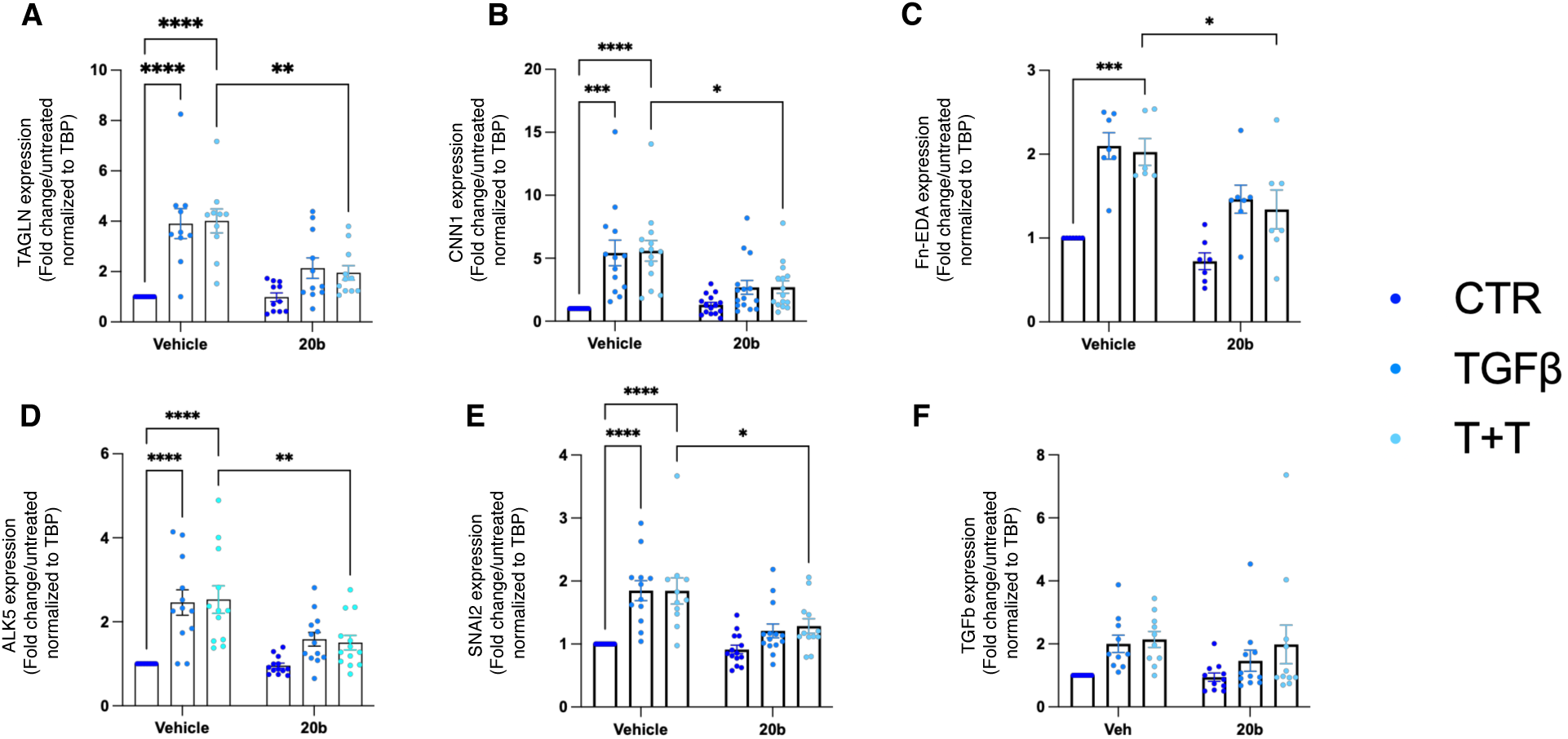
HDAC6 inhibition prevents EndMT after 5 days of treatment. (A) TGLN mRNA expression is induced by TGFβ2 alone or TGFβ2 and thrombin. HDAC6 inhibition significantly reduced TGLN mRNA expression induced by TGFβ2 and thrombin. (N=11; **P<0.01; ****P<0.0001). (B) CNN1 mRNA expression is increased following TGFβ2 alone or TGFβ2 and thrombin. TGFβ2 and thrombin-induced CNN1 mRNA expression by is reduced by TCS20b treatment. (N=15; *P<0.05; ***P<0.001; ****P<0.0001). (C) TGFβ2 and thrombin induced fibronectin-EDA mRNA expression. HDAC6 inhibition reduced fibronectin-EDA mRNA expression following TGFβ1 and thrombin. (N=7; *P<0.05; ***P<0.001). (D) TGFβ2 or TGFβ2 and thrombin increased Alk5 mRNA expression. However, this effect is blocked when HDAC6 activity is inhibited (N=12; **P<0.01; ****P<0.0001). (E) SNAI2 mRNA expression is induced by TGFβ2 alone or TGFβ1 and thrombin. HDAC6 inhibition significantly blocked TGFβ2 and thrombin-induced SNAI2 mRNA expression (N=12; *P<0.05; ***P<0.001; ****P<0.0001). (F) TGFβ2 mRNA expression is not affected by any of the treatment (N=11, ns= not significant).

We also examine endothelial and mesenchymal markers at earlier time point after incubation with TGFβ2 and TGFβ2 with thrombin. At days 2, mRNA expression of TGLN, CNN1, ephrin B2 (EFNB2) and FN1-EDA was induced by TGFβ2 or TGFβ2 and thrombin and blocked by the HDAC6 inhibitor, tubacin A, a selective inhibitor of HDAC6 (Figure 4A). Effects of TGFβ2 or TGFβ2 and thrombin treatment were milder for CNN1 and EFNB2 at day 3. Interestingly, effects of treatments were more pronounced at day 3 compared to day 2 for TGLN and FN1-EDA (Figure 4B). TGFβ mRNA expression was significantly increased by TGFβ2 or TGFβ2 and thrombin and inhibited by tubacin A after 2 days of treatment (Figure 4Ci). However, this effect was not observed after 3 days of treatment confirming what we observed at day 5 (Figure 4Cii). In addition, at days 2 and 3, we observed that ALK5 expression was induced TGFβ2 or TGFβ2 and thrombin and inhibited by tubacin A (Figure 4D). When examining the endothelial markers, eNOS and occludin, their expression were similar in every experimental condition (Supplementary figure 1).

**Figure 4.**
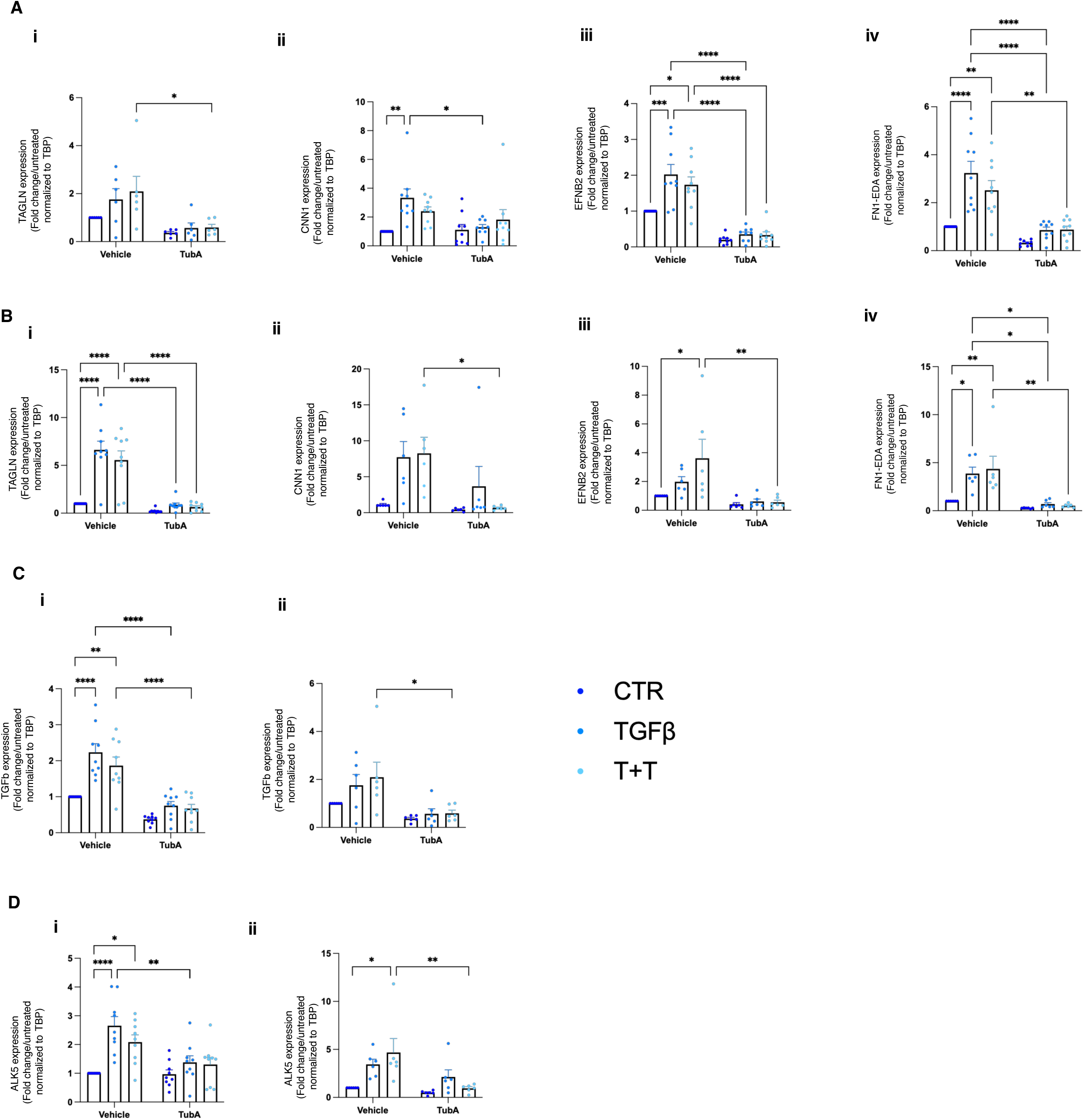
EndMT is prevented by HDAC6 inhibition at earlier time point. (A) mRNA expression of TGLN (i) is not modified by 2 days of TGFβ2 treatment, alone or in combination with thrombin. However, its expression is significantly reduced by HDAC6 inhibition with tubastatin A. Interestingly, mRNA expression of CNN1(ii), EFNB2 (iii) and FN1-EDA (iv) is increased by TGFβ2 and their expression is greatly reduced by HDAC6 inhibition at days 2. (B) After 3 days of treatment, mRNA expression of TAGLN (i) is increased by TGFβ2 treatment, alone or in combination with thrombin. The effect of TGFβ2 or TGFβ2 and thrombin treatment on mRNA expression of CNN1(ii) and EFNB2 (iii) is more modest at day 3 compared to day 2. However, the effect on mRNA expression of FN1-EDA (iv) remains important and dependent on HDAC6 activity. (N=9; *P<0.05; **P<0.01; ***P<0.001; ****P<0.0001).

### HDAC6 mitigates TGFβ signaling

TGFβ2 signal is transmitted into cells through two different pathways: one canonical involving the transcription factors SMADs and one non-canonical mediated through SMAD-independent pathways including ERK1/2 signaling, Rho guanosine triphosphatase (GTPase) signaling, p38 MAPK signaling, JNK signaling, nuclear factor-κB (NFκB) signaling, PI3K/AKT signaling or JAK/STAT signaling. We first evaluated activation of Smad3 in HUVECs stimulated with TGFβ2 and thrombin with or without tubacin A for 30 min and 1h. Smad3 phosphorylation was not modified by any of our experimental conditions (data not shown). Then, we evaluated ERK1/2 activation in HUVECs. ERK1/2 phosphorylation was increased by TGFβ2 treatment. In presence of tubacin A, TGFβ2 no longer induced ERK1/2 phosphorylation (Figure 5).

**Figure 5.**
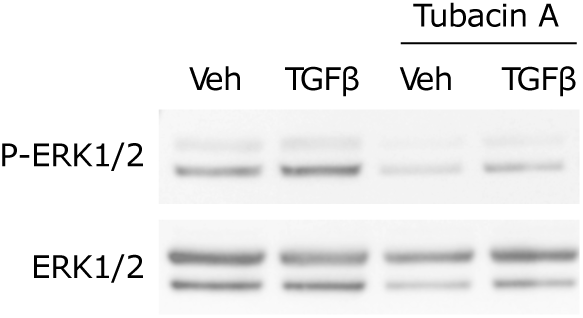
HDAC6 regulates ERK1/2 signaling. Representative western blot images of phosphorylation of ERK1/2 and total ERK1/2 (N=5).

### HDAC6 inhibition delayed thrombus formation and is associated with reduced EndMT

To determine the role of HDAC6 in EndMT in venous thrombosis, we used a mouse model of venous thrombosis where HDAC6 activity was inhibited by treatment with tubastatin A for 7, 14, 17 or 21 days (Figure 6A). Carstairs’ staining revealed significant differences in thrombus size between the tubastatin A groups and the vehicle group at day 7 (Figure 6B). Thrombus size was similar at any point within the tubastatin A group and, as compared to the vehicle group (Figure 6C). These data suggested that inhibition of HDAC6 delayed thrombus formation. Immunofluorescence staining was performed to assess HDAC6 expression. In control and treated mice, HDAC6 expression was observed in the vein wall and the thrombus at each time point studied without any differences between the experimental groups (Figure 6D). Next, we evaluated if venous thrombosis induced EndMT in our model and if HDAC6 was involved in this mechanism. In situ hybridization was performed to quantify TGFβ and TGLN mRNA expression in the thrombus and the vein wall. We observed that TGFβ mRNA expression was more pronounced within thrombi than in the vascular wall (Figure 7A). Plasmatic levels of TGFβ were equivalent between vehicle and treated groups at all time points (Figure 7B). In mice without thrombosis, TGLN mRNA was express in smooth muscle cells underneath the endothelium (Figure 8A). 21 days after thrombosis, in the vehicle-treated group, TGLN mRNA expression appeared in the endothelium. Whereas in tubastatin A-treated animal positive staining for TGLN mRNA was observed underneath the endothelium resembling the condition without thrombosis (Figure 8B).

**Figure 6.**
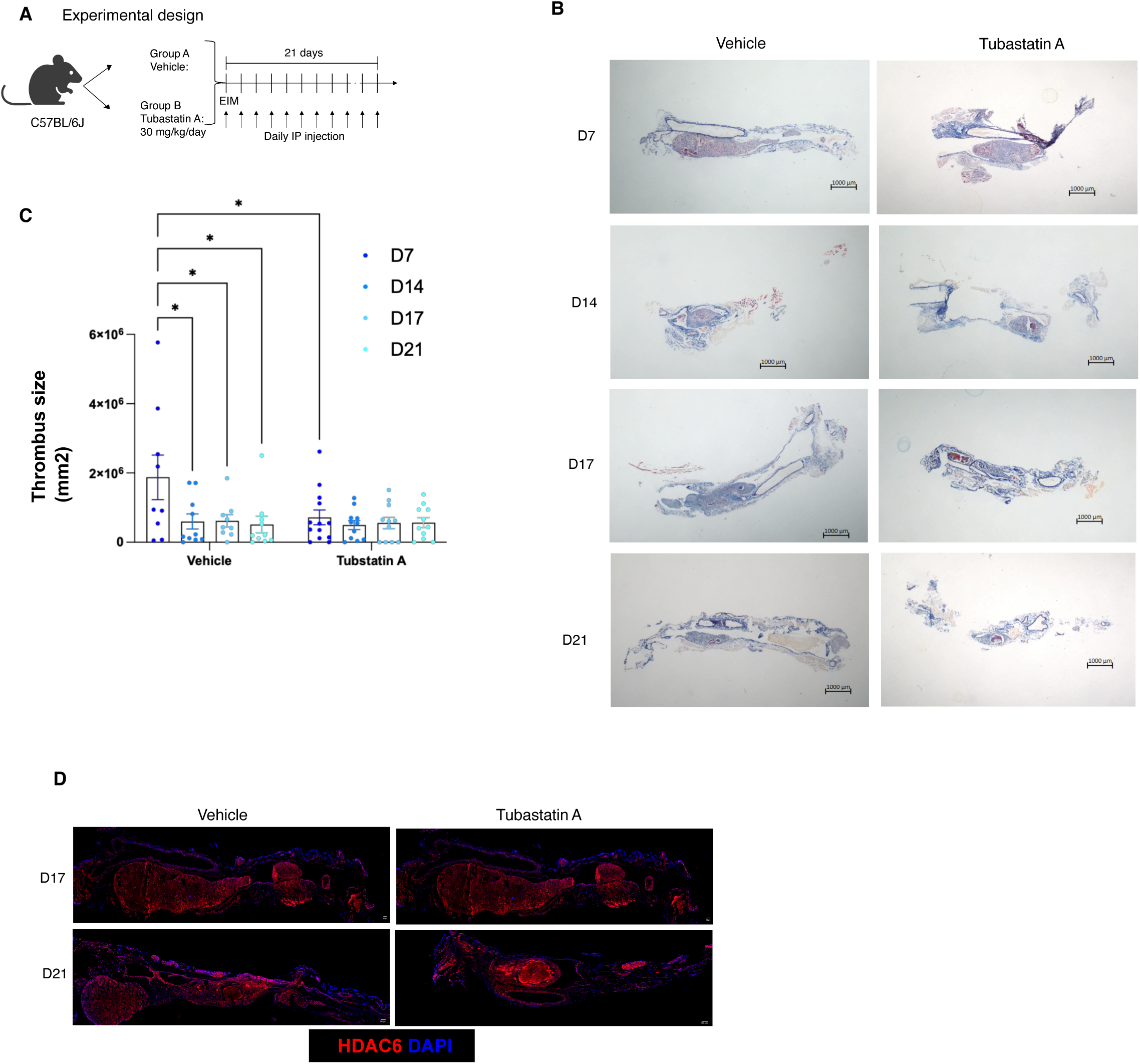
HDAC6 is regulated thrombus formation in vivo. (A) Schematic representation of experimental design for evaluating the effect of HDAC6 inhibition overtime from 0 to 21 days after thrombus induction using the EIM model. (B) Representative images of Carstair’s staining for the 8 experimental groups (Bar=1000 μm). (C) Quantification of Carstair’s staining demonstrated that thrombus size decreases overtime from day 7 to day 21 in the vehicle group. Importantly, thrombus size is significantly smaller in the treated group at day 7 compared to the control group at the same time point. Thrombus size is equivalent at each timepoint when HDAC6 is inhibited (N=11-13; *P<0.05). (D) Representative immunofluorescent staining images of HDAC6 expression in all experimental groups (N=11-13; Bar=200 μm).

**Figure 7.**
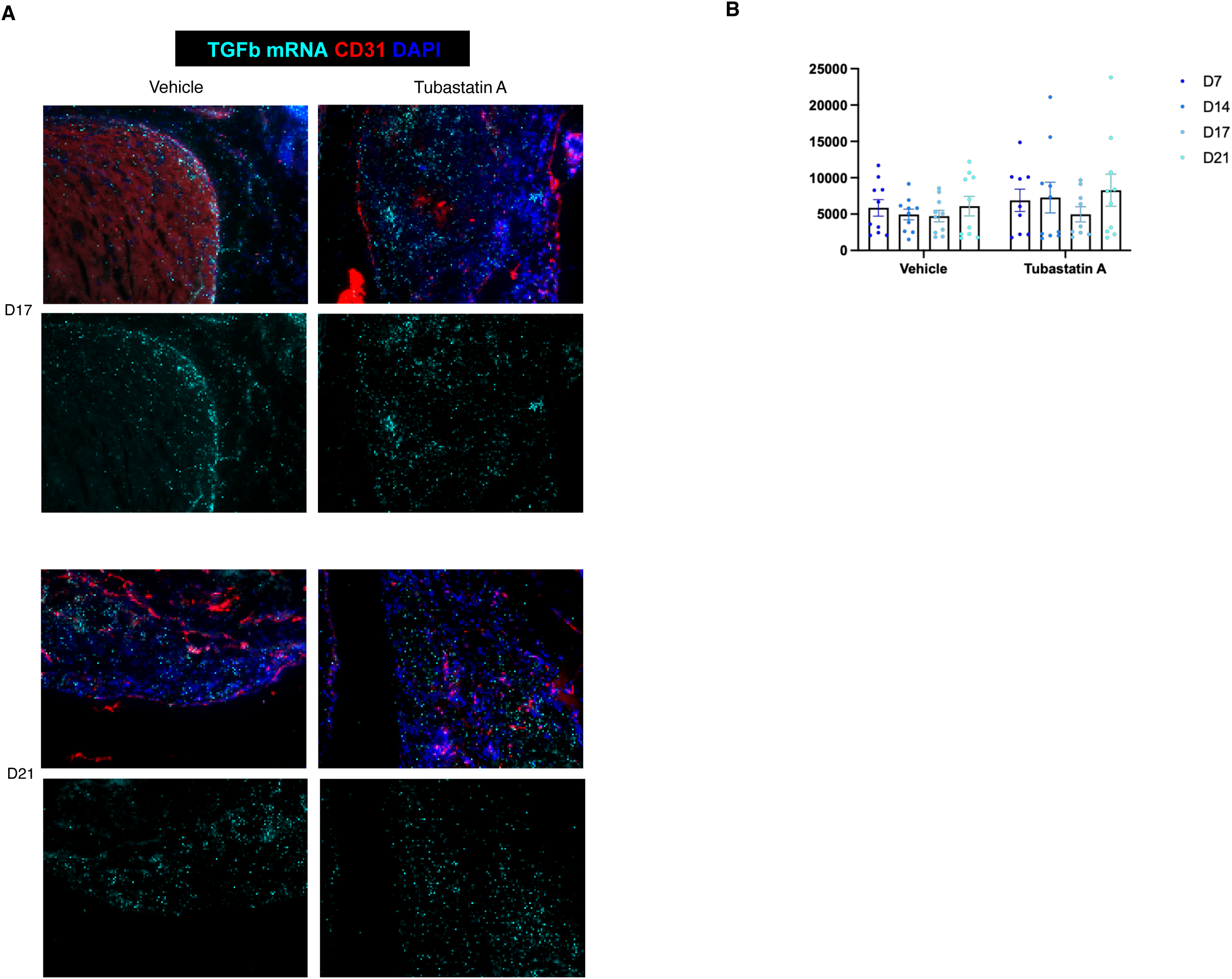
TGFβ expression in thrombus and plasma is not affected by HDAC6 inhibition. (A) mRNA TGFβ transcript within thrombus of vehicle- or HDAC6 inhibitor-treated animals is equivalent as revealed by in situ hybridization (Representative image of 10 independent experiments). (B) Similarly, TGFβ plasmatic levels are not modified by HDAC6 inhibition in mice after venous thrombosis at any time point. (N=10).

**Figure 8.**
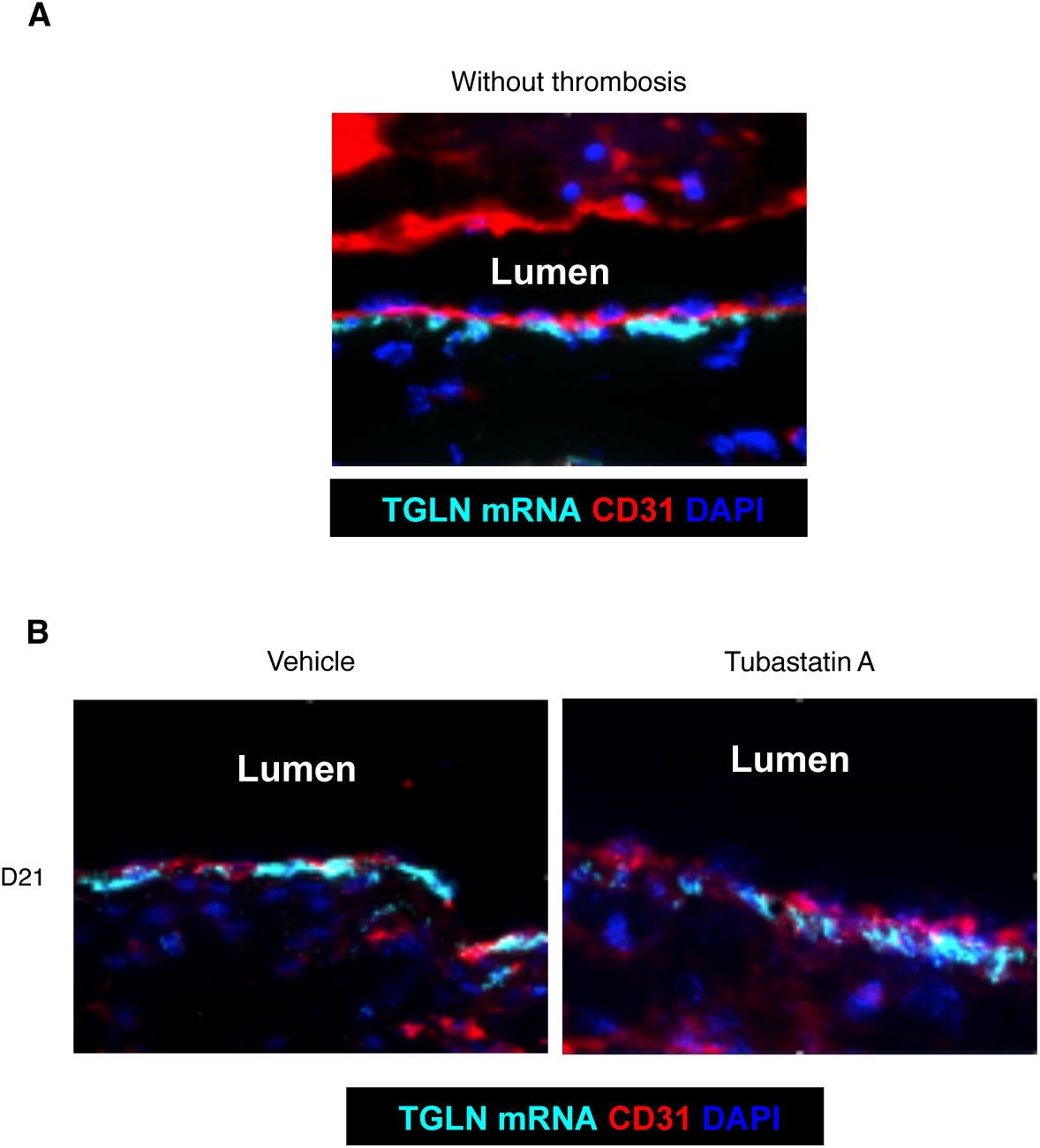
HDAC6 inhibition reduces endothelial expression of SM22 induced by venous thrombosis. (A) Representative immunofluorescent staining images for CD31 (an endothelial marker; red) and TGLN (a mesenchymal marker; light blue) and DAPI-staining nuclei (blue) in samples without thrombosis. (B) At day 21 in the vehicle group, endothelial cells expressed both CD31 and TGLN mRNA. No co-positive cells are detectable in the tubastatin A treated group (Representative image of 10 independent experiments).

We also assessed in the plasma of all experimental groups the levels of the pro-inflammatory cytokines TNFα and IL6 and of the anti-inflammatory cytokines IL10 and IL27. Although none of the statistical comparison reached significance, we observed that TNFα and IL6 tended to be higher in the vehicle groups compared to the tubastatin A-treated animals (Supplementary Figure 2A-B). IL10 was not modified in any of the conditions (Supplementary Figure 2C). On the other hand, IL27 appeared more elevated in the tubastatin A-treated groups, at least at 7, 14 and 17 days of thrombosis (Supplementary Figure 2D). However, further experiments are required to confirm these data.

### Elevated levels of fibronectin-EDA are associated with venous thrombosis

To verify systematic regulation of FN1-EDA under thrombotic conditions, we reanalyzed publicly available RNA-sequencing datasets from two independent studies to investigate the regulation of FN1 isoforms following experimental venous thrombosis. RNA-seq data generated from murine and porcine vein wall and thrombus tissues were systematically analyzed to identify changes in FN1 expression and alternative splicing, with a particular focus on regulation of the EDA-containing isoform during the early response to thrombosis^14,16^.

The RNA-seq analysis revealed species- and vascular bed-specific regulation of FN1-EDA alternative splicing following experimental venous thrombosis (Figure 9). In the murine inferior vena cava ligation model, analysis performed 24h after thrombosis induction demonstrated increased expression of FN1-EDA isoform compared with sham-operated animals, suggesting an early remodeling response of the extracellular matrix following thrombus formation (Figure 9A). In contrast, no significant increase in FN1-EDA expression was detected in the porcine thrombosis model, neither in the femoral vein nor the pulmonary artery at the same time point, suggesting that total FN1-EDA expression may not be uniformly induced across species or thrombosis models (Figure 9B). However, analysis of alternative splicing revealed a distinct tissue-specific pattern in the porcine model, with significantly greater inclusion of the EDA-encoding exon (exon 33) in the femoral vein than in the pulmonary artery under both control and thrombosis conditions (Figure 9C). This finding indicates that regulation of FN1-EDA is governed not only by changes in transcript abundance but also by differential exon inclusion, with the venous environment exhibiting a constitutively higher propensity to generate the EDA-containing isoform.

**Figure 9.**
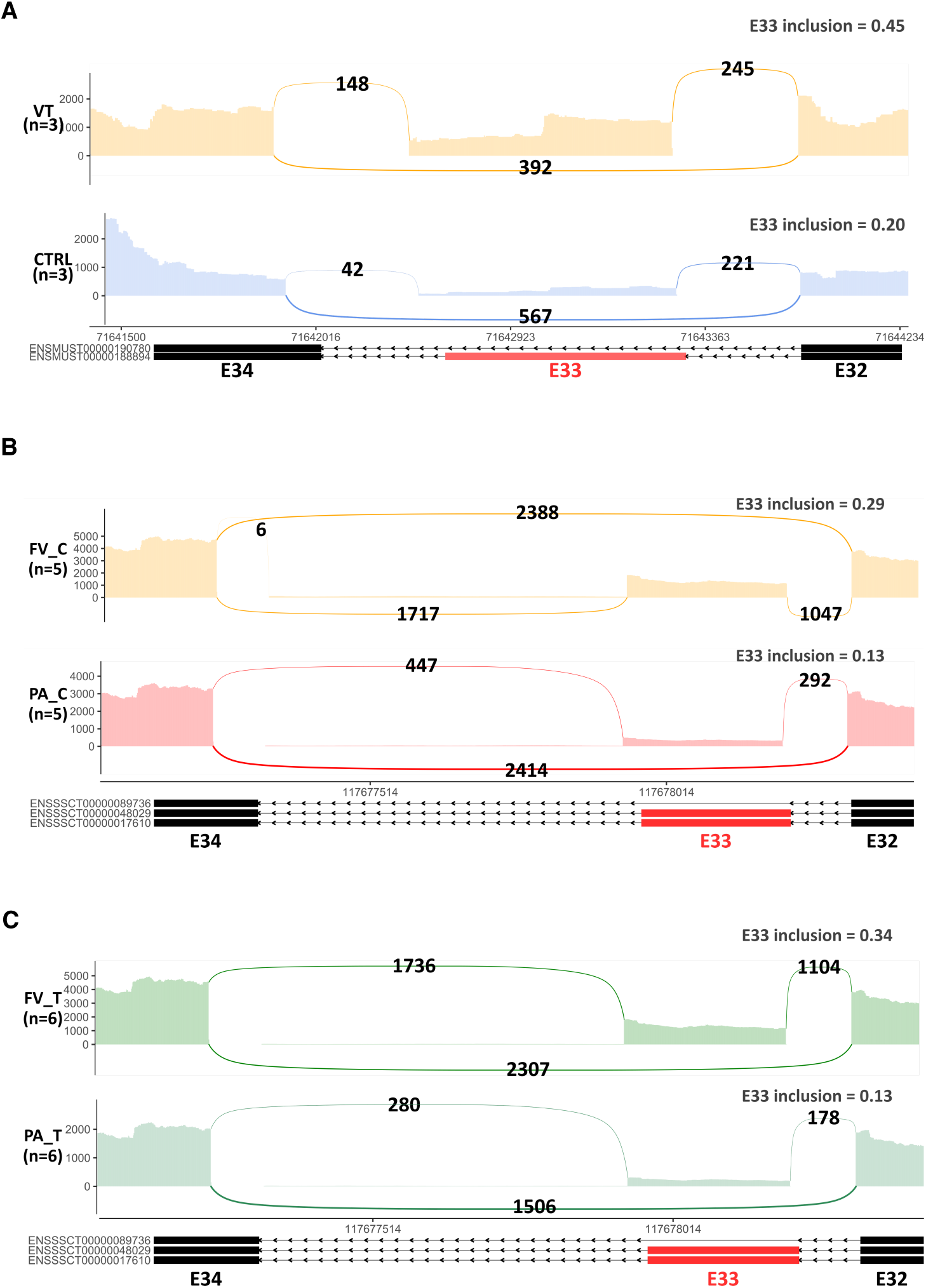
Analysis of the alternative splicing pattern of EDA-coding exon (E33). (A) Sashimiplots reporting the level of expression and inclusion of E33 between murine venous thrombosis models and sham groups (N=3). Read coverage and the number of reads supporting exclusion and inclusion of the EDA-coding exon E33 are given. Comparison of the level of inclusion of E33 between pulmonary artery and femoral veins in the porcine model, in absence of experimental venous thrombosis (B) or under experimental venous thrombosis (C). CTRL: control; FV: femoral vein; FV_C: femoral vein control; FV_T: femoral vein thrombosis; PA: pulmonary artery; PA_C: pulmonary artery control; PA_T: pulmonary artery thrombosis; VT: venous thrombosis.

Based on these data, we measured plasmatic levels of FN1-EDA in patients after a first event of VTE. We selected two groups of patients according to their VTE recurrent status. For both groups, we had plasma sample available after discontinuation of the anticoagulant treatment at 7, 24, 25 and 48 months’ time points. Interestingly, we observed that FN1-EDA levels tended to decrease overtime in patients that did not experience a recurrent DVT event, whereas FN1-EDA levels appeared to remain stable over time in patients with recurrent DVT (Figure 10A). In PE patients, we did not observe any differences of FN1-EDA levels within the recurrent or non-recurrent groups. Although these data are preliminary, we propose that investigating the association between recurrence and FN1-EDA plasmatic levels might represent a new parameter to predict recurrence in patients with idiopathic DVT. Importantly, these data tended to recapitulate our *in vitro* and *in vivo* results.

**Figure 10.**
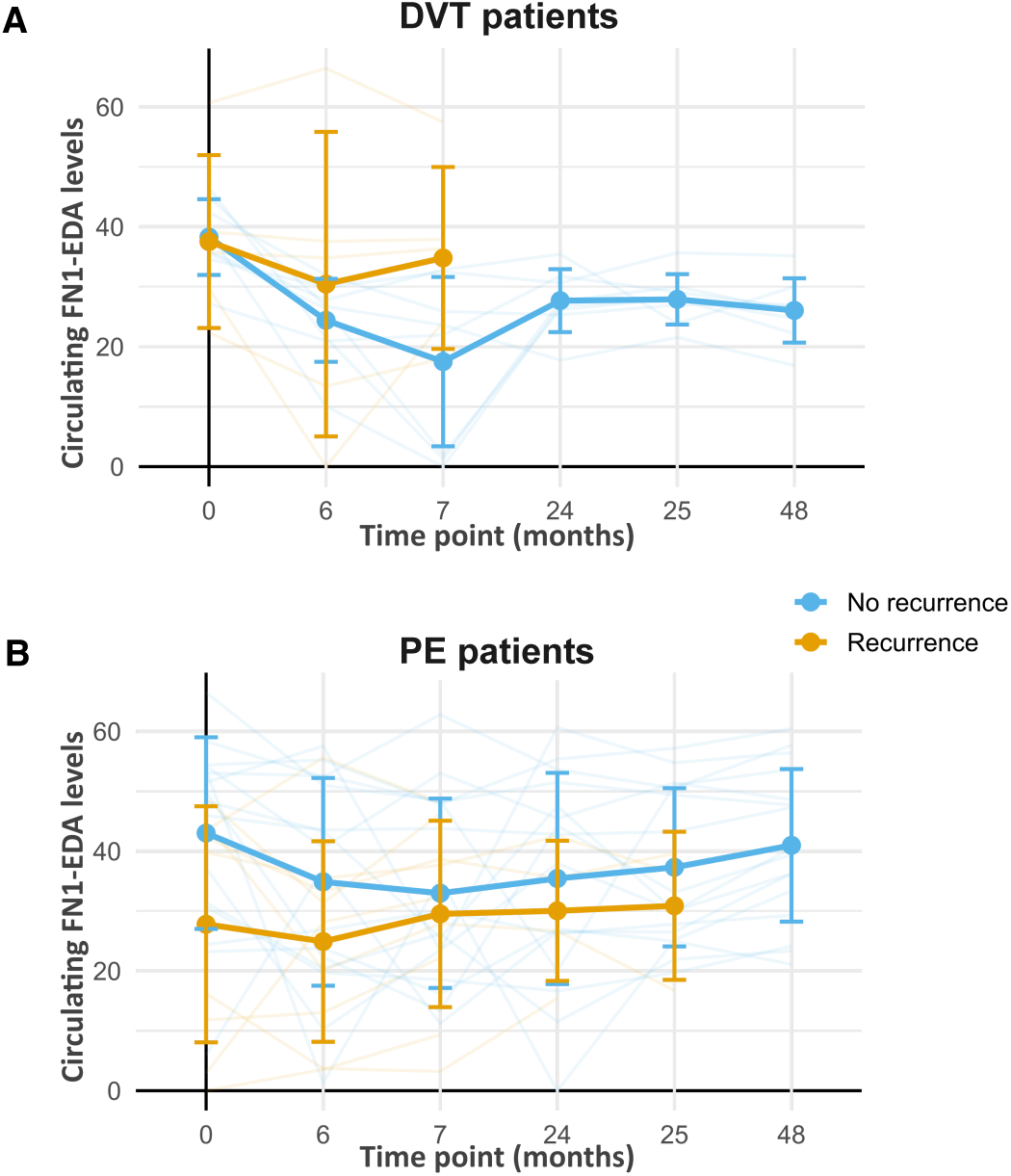
Plasmatic levels of FN1-EDA in patients. (A) Plasmatic levels of FN1-EDA appear higher in patients experiencing recurrent DVT in comparison to non-recurrent patients. Curves of tend to be different around 7 months following the DVT event between patients with recurrent DVT versus non-recurrent patients. This time point corresponds to the plasma samples close to the time of recurrence in these patients. (B) In EP patients, plasmatic levels of FN1-EDA are equivalent in recurrent and non-recurrent patients. (recurrent DVT N=5; non recurrent DVT N=8; recurrent PE N=9; non-recurrent PE N=16).

## Discussion

Current understanding of the mechanisms involved in vein wall fibrosis remains limited and anticoagulant therapy are not helpful to prevent these adverse consequences of VTE. EndMT is a significant process implicated in fibrosis that impairs thrombus resolution. Here, we provide mechanistic insight into the occurrence and the regulation of EndMT following VTE. Previous work has demonstrated that activated TGFβ1 signaling is an important mechanism involved in vascular remodeling following VTE. Hence, TGFβ1, released in large quantity by platelet after venous thrombosis, promotes EndMT through a mechanism involving endothelin-1 overexpression. Importantly, these findings were validated in endarterectomy of chronic thromboembolic pulmonary hypertension (CTEPH) patients ^5,6^. However, the detailed mechanism regulating EndMT after an event of VTE is still unknown.

In the present study, we provide evidence that HDAC6 promotes EndMT and that targeting this pathway might be beneficial to prevent thrombosis-associated EndMT. We found that EndMT occurs in thrombin- and TGFβ2-stimulated HUVECs and following VTE in mice. More specifically, we showed that HDAC6 inhibition was associated with decreased expression of mesenchymal markers and members of the TGFβ2 pathway in endothelial cells. Importantly, we found that HDAC6 inhibition was associated with smaller thrombus size which might reflect enhanced thrombus resolution and less co-positive staining of the endothelial markers CD31 with transgelin suggesting reduced occurrence of EndMT. Finally, we found that FN1-EDA is associated with thrombosis and is a downstream effector of TGFβ-induced HDAC6 activity involved in EndMT.

VTE is associated with long term complications that are thought to contribute to recurrent VTE. Histologically, it is well documented that vascular remodeling following VTE involved fibrosis and accumulation of extracellular matrix. However, how this occurs and what is the trigger is mostly unknown. Venous thrombus resolution resembles wound healing, and several factors have been shown to promote fibrosis and impair thrombus resolution. For example, some human studies have reported that IL6 is a biomarker of post-thrombotic syndrome and venous thrombosis^19^. IL6 is an important mediator of fibrosis and macrophage function during thrombus resolution. IL6 deficiency in mice appears to improve thrombus resolution ^20^. However, in experimental model of venous thrombosis IL6 blocking has failed to improve fibrosis or thrombus resolution^21^. The main source of IL6 during thrombus resolution is the macrophages that contribute to fibrin and matrix degradation. However, the conflicting results obtained following IL6 deficiency or blocking suggest that other cellular mechanisms are in play during thrombus resolution.

TGFβ is also a potent inducer of fibrosis and is thought to greatly contribute to vascular remodeling in the context of thrombosis. The TGFβ family of ligands has been implicated in several deleterious effects during venous thrombosis formation and resolution^22^. Hence, in their recurrent model of VTE in mice, Andraska et al. showed that TGFβ1 and collagen III mRNA expression was increased following two events of thrombosis. This data suggested an association between TGFβ1 and fibrotic remodeling^23^. In line with this study, Bochenek et al. demonstrated that TGFβ signaling delays thrombus resolution and enhances fibrosis in CTEPH patients^24^. They also demonstrated in experimental thrombosis of the inferior vena cava that platelet TGFβ1 deficiency enhances thrombus resolution and, on the contrary, that accumulation of circulating TGFβ1 delays thrombus resolution^25^. Accordingly, we observed high expression of TGFβ within the thrombus in mice using the electrolytic IVC model. TGFβ expression appears to stay stable overtime and do not decrease with thrombus resolution. This is in accordance with the above cited work showing that TGFβ is a major determinant of venous thrombus resolution^25^. Although we did not investigate the source of TGFβ in the present study, there are strong evidence in the literature suggesting that platelets are major contributor. Interestingly, DeRoo et al. showed that platelets contribute to thrombus fibrosis and vein wall remodeling after thrombosis. Platelet depletion decreased collagen content within thrombi without affecting thrombus size. A potential mechanism by which platelet contribute to fibrosis might be by releasing TGFβ release during venous thrombosis^6,26^.

TGFβ is also the most potent inducer of EndMT^27,25,24^. Endothelial cells that undergo EndMT present diminished endothelial properties and enhanced mesenchymal properties including collagen deposition and migration. Thus, EndMT mechanisms have been increasingly studied in several cardiovascular diseases, and it is thought to play an important role in thrombosis-associated vascular remodeling. Accordingly, we found in our *in vitro* conditions that HUVECs undergo EndMT when incubated with TGFβ2 and thrombin. However, in our settings only mesenchymal genes, including transgelin, calponin, ephrin B2 and fibronectin-EDA, and not endothelial genes, were regulated by TGFβ2 and thrombin treatment *in vitro*. Although EndMT has been implicated in venous thrombus resolution, little is known regarding the regulatory pathways involved^28,25^. Interestingly, in different cardiovascular diseases, epigenetic pathways including miRNA and histone or DNA modifications have been shown to regulate EndMT^29^.

We focused on HDAC6, a class IIb HDACs, that is predominantly located in the cytoplasm. It is known to regulate diverse cellular processes including protein aggregation, cytoskeleton rearrangement and cell migration. Importantly, HDAC6 has been involved in EMT in cancer in response to TGFβ1^30^. In the present study, we showed *in vitro* that HDAC6 inhibition led to reduction of mesenchymal gene expression and regulation of components of the TGFβ signaling pathway. These data suggest that HDAC6 might regulate TGFβ intracellular signaling pathways leading to EndMT. In our experimental conditions TGFβ2 did not activate the classical Smad pathway. However, we found that activation of ERK1/2 was dependent on the TGFβ2/HDAC6 axis. Non-canonical TGFβ signaling pathways are involved in multiple cellular function. However, few of them have been linked to EndMT. Studies found that high glucose induced EndMT through regulation of TGFβ and ERK1/2 signaling^31,32^. Hence, inhibition of ERK1/2 has been shown to partially blockade EndMT^31^. Others demonstrated direct activation of ERK1/2 downstream TGFβ-induced EndMT^33^. In atherosclerosis, it was recently showed that ERK1/2 activation is directly associated with EndMT and that its inhibition has beneficial effects on the endothelial phenotype^34^. Accordingly, our data showed that TGFβ mediates its effects on EndMT through activation of HDAC6 and ERK1/2.

In addition to these key findings, our results also demonstrate that HDAC6 inhibition regulates endothelial cell phenotypic changes that were induced following venous thrombosis and led to favorable modulation of thrombus development *in vivo*. Hence, compared to the vehicle treated group, HDAC6 inhibition significantly reduced thrombus size at day 7. This was associated with reduced EndMT as illustrated by co-staining of cells with CD31 and transgelin mRNA probe at day 21. EndMT has been implicated in the impairment of thrombus resolution and fibrosis. TGFβ signaling in endothelial cell contributes to enhanced endothelin-1, which promotes, with TGFβ, extracellular matrix production and fibrosis. Importantly, endothelin-1 receptor blocking prevented gene expression of transgelin and collagen and restored thrombus resolution^6^. Although, it would require further investigation in the context of venous thrombosis and EndMT, some data from the literature suggest that endothelin-1 expression might be regulated by HDAC6^35^.

Importantly, our analyses identified proteins induced by TGFβ and regulated by HDAC6, at least *in vitro*, in endothelial cells. Ephrin B2 and fibronectin-EDA were significantly increased by TGFβ and regulated by HDAC6. Interestingly, ephrin B2 has been involved in EndMT in arterial endothelial cells in diabetic conditions. Silencing of ephrin B2 or downstream signaling pathway FAK partially blocked EndMT in response to high glucose^36^. Furthermore, ephrin B2 has been shown to promote endothelial cell migration in HUVECs^37^. This result supports the role of HDAC6 in functionally regulating EndMT in response to TGFβ. FN1-EDA is a marker consistently regulated by HDAC6 in our *in vitro* experiments.

FN1 has been involved both in EndMT and thrombo-inflammatory processes^38,39^. It is one of the gene and encoded protein that is induced during EndMT, characteristic of a mesenchymal phenotype. FN1 is a dimeric extracellular matrix protein including different type of repeating segments, type I, type II and type III, which are constitutively expressed. It also includes 2 segments, extra domain A (EDA) and extra domain B (EDB) that are alternatively spliced. This results in two major isoforms: plasma FN1, secreted by hepatocyte which lacks both EDA and EDB and cellular FN, produced by cells and with variable EDA or EDB segment inclusion. FN1 is a glycoprotein involved in hemostasis and in clot stabilization and is upregulated in mouse in a model of iliac compression^7^. In pathological conditions such as stroke, FN1-EDA plasmatic levels are increased and promote arterial thrombosis through TLR4 signaling^40^. Importantly, the presence of FN1-EDA is characteristic of tissue repair and fibrosis. FN1-EDA deficient mice are protected against bleomycin-induced lung fibrosis and fibroblasts exert reduced capacities to respond to TGFβ^41^. FN1-EDA has been found to bind and recruit latent tumor growth factor TGFβ1 binding protein (LTBP) to the extracellular matrix, which promote the sequestration and the activation of TGFβ1^42^. Here, we provide evidence that FN1-EDA is upregulated as soon as 48h in HUVECs treated with TGFβ in a HDAC6-dependent manner. We analyzed publicly available RNAseq data sets and showed that expression of FN1-EDA was increased in a murine model of thrombosis compared to sham operated animals. This regulation was specific to the inferior vena cava as FN1-EDA expression was not associated with PE. Importantly, this analysis validates our data in an independent data set.

Additionally, FN1-EDA plasmatic levels are increased in overweight patient with VTE compared to control^39^. Based on our experimental data, we measured FN1-EDA in cohorts of patients with DVT or PE with a normal BMI and assess their plasmatic level evolution in time in the same patient. When we classified patients according to their recurrent status, we observed that plasma levels of FN1-EDA were decreasing overtime in non-recurrent DVT patients. On the contrary, in recurrent DVT patients, FN1-EDA plasma levels were stable overtime. Our data suggest that lack of FN1-EDA downregulation might favor recurrent DVT. In PE patients, levels of FN1-EDA remained similar regardless of the recurrent status. The exact mechanism by which FN1-EDA is involved in recurrent DVT would require further investigation. However, one can hypothesize that a defect in the regulation of FN1-EDA might be associated with higher risk of recurrence. These data might have important implications for the management of recurrent thrombosis since FN1-EDA has been involved in fibrosis. Hence, one can speculate that enhancing thrombus resolution and preventing fibrosis might improve vascular physiology and provide better parameter to stratify patients according to their risk of recurrence.

Collectively our data suggest that targeting HDAC6 might represent an attractive therapeutic strategy. HDAC6 inhibitors exert low toxicity compared to general inhibitor of HDACs because of its non-histone function. This may be illustrated by the use of valproic acid in an experimental model of thrombosis. Hence, it appears that valproic acid significantly reduced thrombus size when used at low dose whereas at higher dose it exerts pro-thrombotic properties^43^. This study showed that there is a need for more targeted strategies. However, tubastatin A, which is the most widely used and most specific HDAC6 inhibitor, has low biodisponibility. Other molecules including rocilinostat (ACY-1215) and citarinostat (ACY-241) might represent better strategies. Importantly, HDAC6 inhibitors are safe and present advantages compared to TGFβ inhibition. Some of these compounds have been used in lung fibrosis and have shown promising effects^44^.

Our data showed for the first time that HDAC6 has a unique role during venous thrombosis by promoting EndMT, which is associated with increased thrombus burden. We have demonstrated that selective inhibition of HDAC6 is effective in reducing EndMT *in vitro* and *in vivo* suggesting that targeting HDAC6 might be beneficial to prevent vascular damages associated with thrombosis that might be involved in recurrent thrombosis. Importantly, our data also suggest that FN1-EDA might be an important biomarker related to VTE recurrence. Our work also provides solid ground for the evaluation of this biomarker in VTE patients to help stratify their risk of recurrence.

## Author contributions

M. Pilard performed research, analyzed data and wrote the manuscript. V. Gourdou-Latyszenok, L. Civi, S. Robin, L. Gourhant, E. L. Ollivier, T. Riou, J. Elhasnaoui, L. Bicrel, R. Ricci de Azevedo, G. Pernod performed research and analyzed data. C. Tromeur and F. Couturaud interpreted the data and wrote the manuscript. C. A. Lemarié conceived and supervised the study, performed research, analyzed data and wrote the manuscript.

## Acknowledgment

This study was supported by Fondation du Souffle (M.P.). C.A.L. was supported by Région Bretagne, Département du Finistère and Brest Métropole. C.A.L. also received funding from the “Fédération Française de Cardiologie” and “Le Nouveau Souffle”. We would like to acknowledge and thank the animal care facility for their technical assistance.

