## Supplemental files for "HDAC6 is a novel regulator of endothelial-to-mesenchymal transition in venous thrombosis"

**Supplementary data**

**List of PADIS-PE and PADIS-DVT investigators**

- PADIS-PE Investigators:

Francis Couturaud[1], Olivier Sanchez[2], Gilles Pernod[3], Patrick Mismetti[4], Patrick Jego[5], Elisabeth Duhamel[6], Karine Provost[7], Claire Bal dit Sollier[8], Emilie Presles[9], Philippe Castellant[10], Florence Parent[11], Pierre-Yves Salaun[12], Luc Bressollette[13], Michel Nonent[14], Philippe Lorillon[15], Philippe Girard[16], Karine Lacut[1], Marie Guégan[1], Jean-Luc Bosson[17], Silvy Laporte[9], Christophe Leroyer[1], Hervé Décousus[4], Guy Meyer[2], Dominique Mottier[1].

Affiliations
[1] Département de Médecine Interne et Pneumologie, Centre Hospitalo-Universitaire de Brest, Université de Bretagne Occidentale, and EA 3878, CIC INSERM 1412, Brest, France.
[2] Groupe d'Investigation et de Recherche Clinique sur la Thrombose (GIRC Thrombose). Service de Pneumologie, Hôpital Européen Georges Pompidou, AP-HP; Université Paris Descartes, Sorbonne Paris Cité, and INSERM UMR S 970, Paris, France.
[3] Groupe d'Investigation et de Recherche Clinique sur la Thrombose (GIRC Thrombose). Département de Médecine Vasculaire, Centre Hospitalo-Universitaire de Grenoble, Université de Grenoble 1, Grenoble, France.
[4] Groupe d'Investigation et de Recherche Clinique sur la Thrombose (GIRC Thrombose). Service de Médecine et Thérapeutique, Unité de Pharmacologie Clinique, Centre Hospitalo-Universitaire de Saint-Etienne, and EA3065, Université Jean Monnet, Saint-Etienne, Fr.
[5] Groupe d'Investigation et de Recherche Clinique sur la Thrombose (GIRC Thrombose). Service de Médecine Interne, Centre Hospitalo-Universitaire de Rennes, Université de Rennes 1, Rennes, France.
[6] Groupe d'Investigation et de Recherche Clinique sur la Thrombose (GIRC Thrombose). Service de Médecine Interne, Centre Hospitalier Général de Saint-Brieuc, Saint-Brieuc, France.
[7] Service de Cardiologie, Centre Hospitalier Général de Lannion, Lannion, France.
[8] Clinique des Anticoagulants d'Ile de France (C.R.E.A.T.I.F.), Centre Hospitalo-Universitaire de Lariboisière, Paris, France.
[9] Groupe d'Investigation et de Recherche Clinique sur la Thrombose (GIRC Thrombose). Unité de recherche clinique, Innovation et pharmacologie, Centre Hospitalo-Universitaire de Saint-Etienne, and EA3065, Université Jean Monnet, Saint-Etienne, France.
[10] Service de Cardiologie and EA 4324, Centre Hospitalo-Universitaire de Brest, Université de Bretagne Occidentale, Brest, France.
[11] Groupe d'Investigation et de Recherche Clinique sur la Thrombose (GIRC Thrombose). Service de Pneumologie and INSERM 999, Centre Hospitalo-Universitaire de Kremlin Bicêtre, Kremlin Bicêtre, France.
[12] Groupe d'Investigation et de Recherche Clinique sur la Thrombose (GIRC Thrombose). Service de Médecine Nucléaire and EA 3878, Centre Hospitalo-Universitaire de Brest, Université de Bretagne Occidentale, Brest France.
[13] Groupe d'Investigation et de Recherche Clinique sur la Thrombose (GIRC Thrombose). Service d'Echo-doppler Vasculaire, and EA 3878, CIC INSERM 1412, Centre Hospitalo-Universitaire de Brest, Université de Bretagne Occidentale, Brest, France.
[14] Groupe d'Investigation et de Recherche Clinique sur la Thrombose (GIRC Thrombose). Service de Radiologie, and EA 3878, CIC INSERM 1412, Centre Hospitalo-Universitaire de Brest, Université de Bretagne Occidentale, Brest, France.

[15] Pharmacie Centrale, Centre Hospitalo-Universitaire de Brest, Université de Bretagne Occidentale, Brest, France.
[16] Groupe d'Investigation et de Recherche Clinique sur la Thrombose (GIRC Thrombose)17Département Thoracique, Institut Mutualiste Montsouris; Paris, France.
[17] Groupe d'Investigation et de Recherche Clinique sur la Thrombose (GIRC Thrombose)18Centre d'Investigation Clinique and UMR CNRS 5525, Centre Hospitalo-Universitaire de Grenoble, Université de Grenoble 1, Grenoble, France.

- PADIS-DVT Investigators:

Francis Couturaud[1], Gilles Pernod[2], Emilie Presles[3], Elisabeth Duhamel[4], Patrick Jego[5], Karine Provost[6], Brigitte Pan-Petesch[7], Claire Bal Dit Sollier[8], Cécile Tromeur[9], Clément Hoffmann[10], Luc Bressollette[10], Philippe Lorillon[11], Philippe Girard[12], Emmanuelle Le Moigne[9], Aurelia Le Hir[9], Marie Guégan[9], Silvy Laporte[3], Patrick Mismetti[13], Karine Lacut[9], Jean-Luc Bosson[14], Laurent Bertoletti[13], Oliver Sanchez[14], Guy Meyer[14], Christophe Leroyer[9], Dominique Mottier[9].

Affiliations
[1] Départementde Médecine Interne et Pneumologie, CHU de Brest, Université de Bretagne Occidentale, EA 3878, CIC INSERM 1412, F-CRIN INNOVTE, Brest.
[2] Département de Médecine Vasculaire, CHU de Grenoble, Université de Grenoble 1, F-CRIN INNOVTE, Grenoble.
[3] Unité de Recherche Clinique, Innovation et Pharmacologie, CHU de Saint-Etienne, and INSERM U1059 SAINBIOSE, Université Jean Monnet, F-CRIN INNOVTE, Saint-Etienne.
[4] Service de Médecine Interne, Centre Hospitalier Général de Saint-Brieuc, F-CRIN INNOVTE, Saint-Brieuc.
[5] Service de Médecine Interne, CHU de Rennes, Université de Rennes 1, Rennes.
[6] Service de Cardiologie, Centre Hospitalier Général de Lannion, Lannion.
[7] Service d'Hématologie, Centre Hospitalier Général de Quimper, Quimper.
[8] Clinique des Anticoagulants d'Ile de France (C.R.E.A.T.I.F.), CHU de Lariboisière, Paris.
[9] Départementde Médecine Interne et Pneumologie, CHU de Brest, Université de Bretagne Occidentale, EA 3878, CIC INSERM 1412, F-CRIN INNOVTE, Brest.
[10] Service d'Echo-Doppler Vasculaire, and EA 3878, CIC INSERM 1412, CHU de Brest, Université de Bretagne Occidentale, F-CRIN INNOVTE, Brest.
[11] Pharmacie Centrale, CHU de Brest, Université de Bretagne Occidentale, Brest.
[12] Département Thoracique, Institut Mutualiste Montsouris, F-CRIN INNOVTE, Paris.
[13] Service de Médecine Vasculaire et Thérapeutique, Unité de Pharmacologie Clinique, CIC1408, CHU de Saint-Etienne, and INSERM U1059 SAINBIOSE, Université Jean Monnet, F-CRIN INNOVTE, Saint-Etienne.
[14] CIC and UMR CNRS 5525, CHU de Grenoble, Université de Grenoble 1, Grenoble, France.

**Supplementary data figure legend**

**Supplementary Figure 1.** Endothelial markers are not modified by TGFβ2 or HDAC6 inhibition. (A) mRNA expression of eNOS is not modified by 2 (i) or 3 (ii) days of TGFβ2 treatment, alone or in combination with thrombin. HDAC6 inhibition with tubastatin A had no effect on eNOS mRNA expression. (B) mRNA expression of occludin is not modified by 2 (i) or 3 (ii) days of TGFβ2 treatment, alone or in combination with thrombin. HDAC6 inhibition with tubastatin A had no effect on eNOS mRNA expression (N=8).

**Supplementary Figure 2.** Plasmatic levels of pro- and anti-inflammatory cytokines after venous thrombosis. Plasmatic levels of TNFα (A), IL6 (B), IL10 (C) and IL27 (D) are not modified overtime following venous thrombosis in mice. In the experimental group where HDAC6 was inhibited similar levels of these molecules were quantified. However, TNFα, IL6 and IL27 tended to be increased compared to the tubastatin A group (N=10).

**Table I.** Characteristics of the overall population study

| | Overall (N=38) |
|---|---|
| Age, years | 57.5 (43-66) |
| Female gender | 15 (39.5) |
| Body mass index, kg.m$^{-2}$ | 25.4 (23.5-28.0) |
| VTE recurrence | 14 (37) |
| DVT | 5 (38) |
| PE | 9 (36) |

Continuous variables are summarized as medians (interquartile range: IQR), categorical variables are reported as absolute frequencies (percentages). Abbreviations: VTE, venous thromboembolism; DVT, deep vein thrombosis and PE, pulmonary embolism.

Supplementary Figure 1.

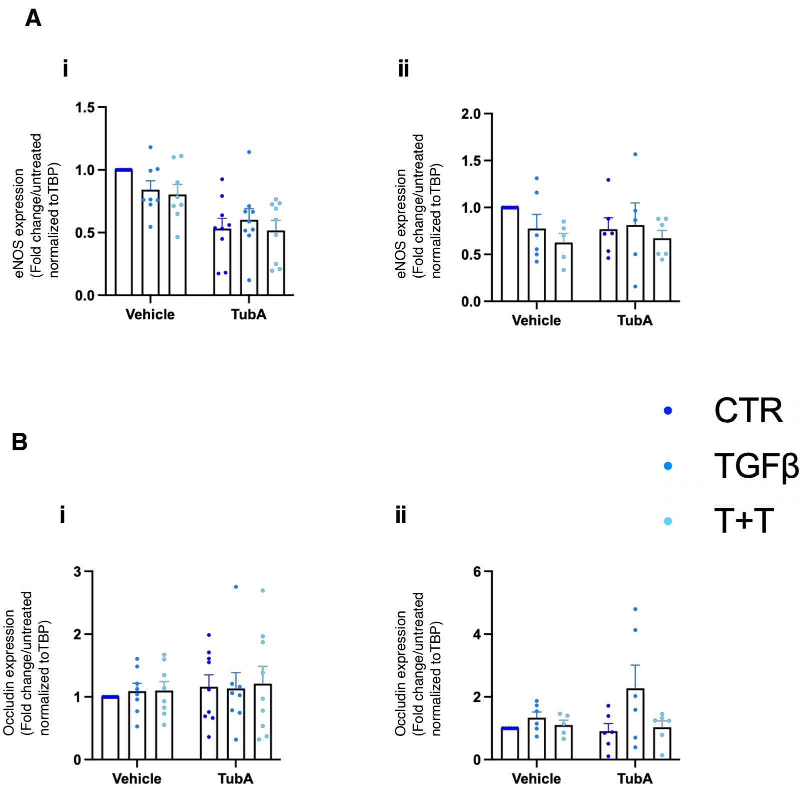

### Supplementary Figure 2

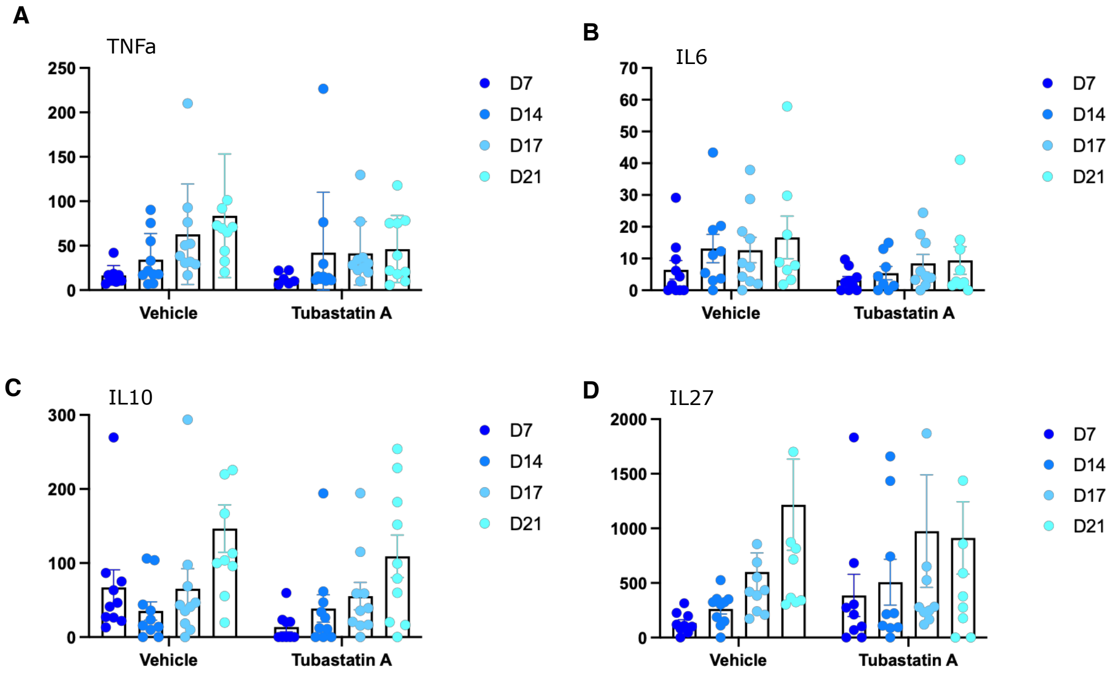
